# Viral surveillance of invasive mammals in New Zealand reveals unique viral lineages reflecting their introduction history

**DOI:** 10.1101/2025.08.27.672626

**Authors:** Rebecca K. French, Florian Pichlmueller, Stephanie J. Waller, Jeremy Dubrulle, Jess Tuxford, Andrew Veale, Jemma L. Geoghegan

## Abstract

Introduced mammalian species in Aotearoa New Zealand pose significant ecological risks and may serve as reservoirs for novel or emerging infectious diseases. In this study, we present the first metatranscriptomic survey of viruses in five introduced mammals: ferrets (*Mustela furo*), stoats (*Mustela erminea*), weasels (*Mustela nivalis*), brushtail possums (*Trichosurus vulpecula*), and European hedgehogs (*Erinaceus europaeus*), sampled across both the North and South Islands. Through total RNA sequencing, we identified 11 mammalian-infecting viruses spanning eight viral families, including four novel virus species: *Ferret mastadenovirus, Possum astrovirus, Ferret pestivirus,* and *Weasel jeilongvirus*. Whole genomes were recovered for six of these viruses, enabling detailed phylogenetic analysis. Notably, we observed strong global geographic clustering in both *Wobbly possum disease virus* and *Ferret hepatitis E virus*, suggesting localized viral evolution following the introduction of their hosts into New Zealand. In addition, the detection of *Human rotavirus A* in hedgehogs highlights the possibility of reverse zoonotic transmission. Together, these findings broaden our understanding of the viral diversity harboured by New Zealand’s introduced mammals and provide a critical foundation for future biocontrol and disease surveillance efforts.

**IMPORTANCE:** Introduced mammals in Aotearoa New Zealand not only threaten native biodiversity through predation and competition, but also represent a largely overlooked source of infectious disease risk. Viruses circulating in these species may spill over into native wildlife, livestock, or even humans, while human viruses can also establish in introduced animals and create new reservoirs. Understanding which viruses are present, and how they evolve in isolated host populations, is critical for anticipating future disease outbreaks, improving biosecurity, and guiding wildlife management strategies. This work provides foundational knowledge that links ecology, conservation, and health, highlighting the need to consider pathogens as part of the broader impact of invasive species.

## INTRODUCTION

Wild mammals are key reservoirs of zoonotic pathogens, posing significant risks to human health (1). These animals harbour a wide range of zoonotic and pathogenic viruses including rabies virus (2), hantavirus (3, 4), morbillivirus (5) and coronaviruses (6). Due to their role in disease ecology, wild mammals have become a central focus of virological surveys, where hundreds of novel viruses have been uncovered across many hosts including in bats, rodents and shrews (7–9). These investigations have often identified viruses closely related to those that cause disease in humans or other animals (8), as well as uncovering novel viruses, many with zoonotic potential (1).

Largely isolated from the rest of the world for more than 50 million years, Aotearoa New Zealand has a unique ecosystem with only two species of bats as the only extant native terrestrial mammals (10). However, numerous mammalian species have been introduced into New Zealand, mainly by European settlers since the 19^th^ century, including stoats, weasels, ferrets, hedgehogs and possums (11). Invasions of mammalian species are one of the major causes of global biodiversity loss and ecosystem change (12), and in New Zealand these mammalian pests have been responsible for both local and country-wide extinctions of many native animals due to predation (13). These mammals are also likely to harbour numerous viruses, either introduced via their native ranges or those that have emerged since their introduction. It is therefore likely that mammalian pests are reservoirs for a vast array of viruses, potentially posing a significant threat to endemic species with no prior exposure (14), as well as to public health.

Very little is known about viruses in New Zealand mammals, with the primary focus for viral surveillance being limited to domestic animals. Viruses have been identified in New Zealand’s bats (15, 16), sea lions (17), long-finned pilot whales (*Globicephala melas*) (18), fur seals (*Arctocephalus forsteri*) (19) and Maui dolphins (*Cephalorhynchus hectori maui*) (20). Pathogenic viruses have also been identified in some wild introduced species including *Rabbit hemorrhagic disease virus* in rabbits (21) and *Wobbly possum disease virus* in brushtail possums (22), as well as serological evidence that stoats are infected with multiple viruses (23). Nevertheless, the total assemblage of viruses (i.e. the virome) of many mammalian pests in New Zealand remains unexplored.

Herein, we used total RNA sequencing to uncover the viruses harboured in five introduced mammals in New Zealand, including ferrets (*Mustela furo*), stoats (*Mustela erminea*), weasels (*Mustela nivalis*), brushtail possums (*Trichosurus vulpecula*) and European hedgehogs (*Erinaceus europaeus*), sampled across both the North and South Islands. Since ferrets, stoats and weasels belong to the *Mustela* genus (Fig. 1A) and geographically overlap (24), we hypothesise that there is frequent cross-species virus transmission between these species compared to brushtail possums and hedgehogs. As the first virome-scale survey of introduced mammals in New Zealand, this study offers new insights into the evolutionary relationships and geographic distribution of mammalian viruses.

**FIG 1.**
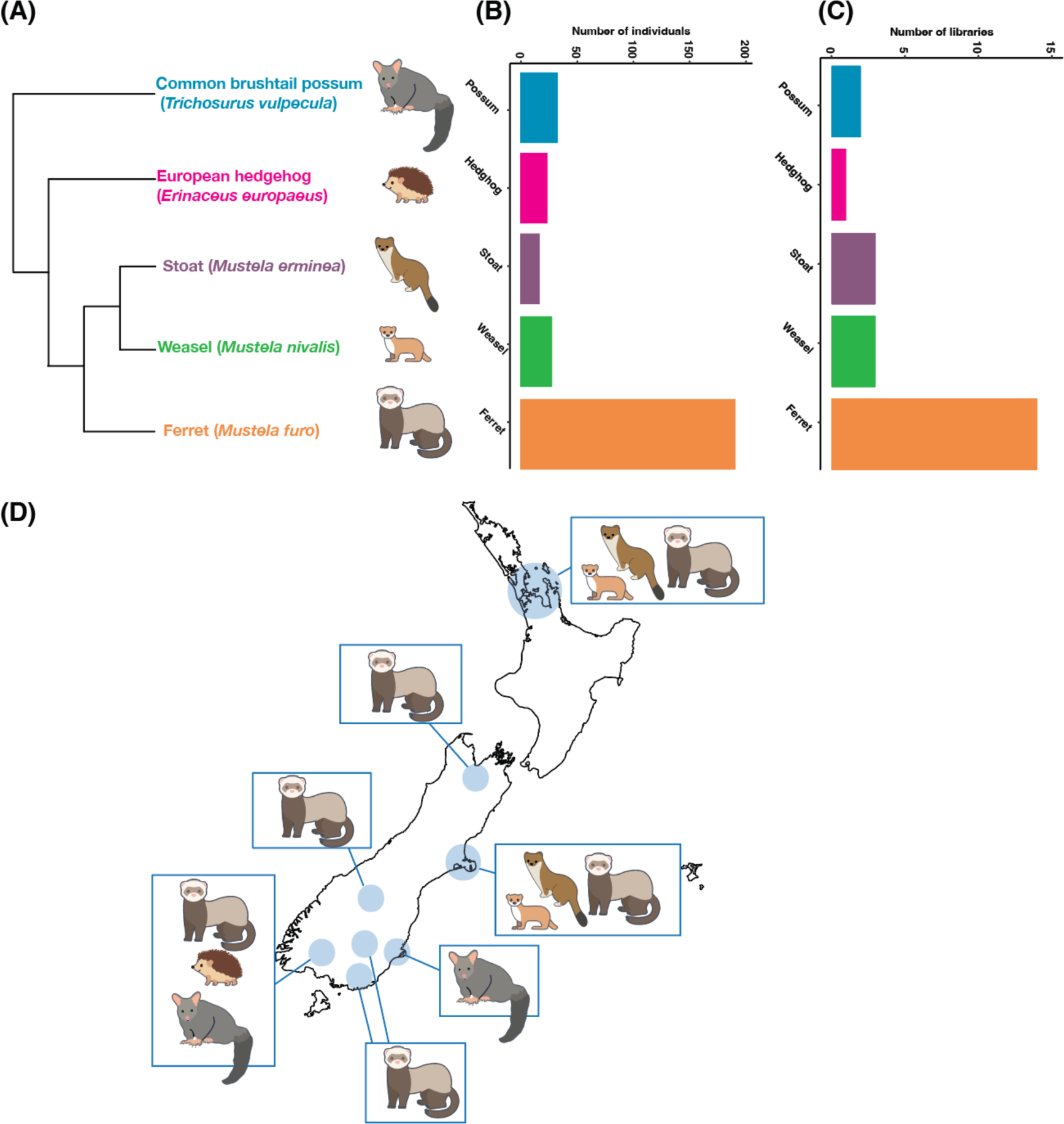
(A) Phylogeny of sampled mammals (based on the Open Tree of Life https://tree.opentreeoflife.org/). (B) The number of individual animals sampled per species. (C) The number of RNA-Seq libraries generated per species. (**D**) Map of New Zealand showing the sampling locations for each species.

## RESULTS

We generated a total of 1.6 billion sequencing reads, with an average of 68 million per library (±11.2 million SD, Fig. 2). The hedgehog library had the highest viral abundance (>36,000 RPM, Fig. 2, Table S2), while other libraries with high viral abundance included the two possum libraries (>800 RPM, Fig. 2, Table S2) and one ferret library (>1,500 RPM, Table S2). No viruses were detected in stoats sampled here.

**FIG 2.**
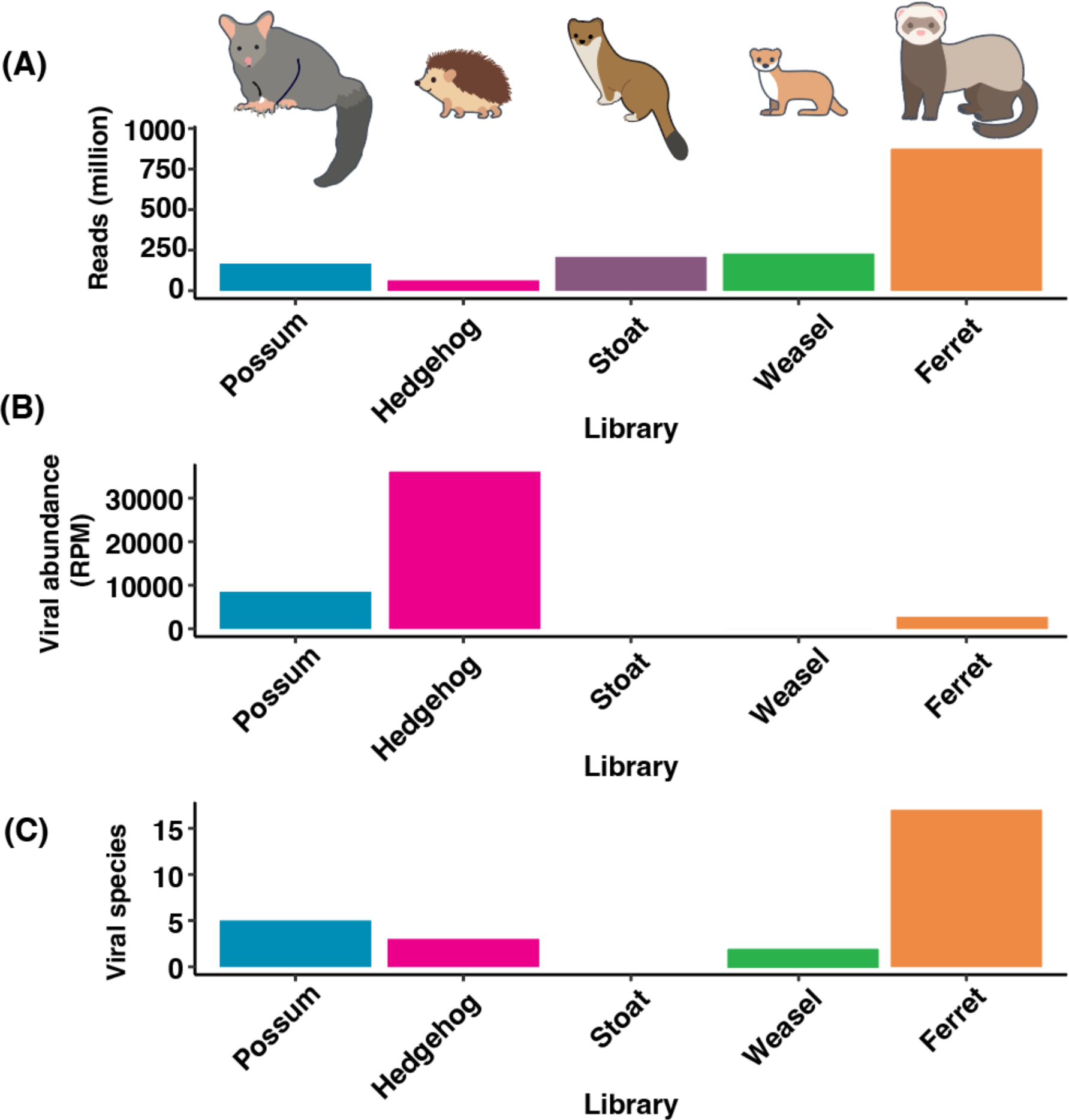
(A) The total number of reads (expressed in millions) for each host species sampled. (B) The total viral abundance (expressed in reads per million, RPM) per host species. (C) The number of viral species detected per host species.

We identified 20 virus transcripts that are known to infect mammals spanning 11 virus species from eight viral families (Table 1), varying in abundance from 1 RPM (*Weasel jeilongvirus*) to >35,000 RPM (*Hedgehog hepatovirus*). Of these 11 virus species, four were considered novel since they shared <90% amino acid sequence similarity to other known viruses within the most conserved region (i.e. the polymerase). We recovered complete viral genomes from six virus species. We now describe the different groups of viruses in turn.

**Table 1.** Details of the viruses found in each RNA sequencing library. Abundance is expressed in reads per million, and the length in nucleotides. CDS = coding sequence. RdRp = RNA dependent RNA polymerase.

| ORGANISM | GENBANK<br>ACCESSION | HOST(S) | LIBRARY | VIRAL FAMILY | GENOME<br>TYPE | ABUNDANCE | NOVEL | GENE(S) | LENGTH |
| --- | --- | --- | --- | --- | --- | --- | --- | --- | --- |
| FERRET MASTADENOVIRUS |  | Ferret | M1 | *Adenoviridae* | ds DNA | 4.8 | Yes | DNA polymerase, partial<br>CDS | 936 |
| HEDGEHOG ARTERIVIRUS |  | Hedgehog | M24 | *Arteriviridae* | + ss RNA | 219 | No | ORF1ab polyprotein,<br>partial CDS | 3931 |
| WOBBLY POSSUM DISEASE<br>VIRUS |  | Possum | M22 | *Arteriviridae* | + ss RNA | 348 | No | Whole genome | 13306 |
| WOBBLY POSSUM DISEASE<br>VIRUS |  | Possum | M23 | *Arteriviridae* | + ss RNA | 839 | No | Whole genome | 13305 |
| POSSUM ASTROVIRUS |  | Possum | M22 | *Astroviridae* | + ss RNA | 52 | Yes | ORF1a Protein, ORF1b<br>Protein, ORF2 Protein<br>partial CDS | 5446 |
| POSSUM ASTROVIRUS |  | Possum | M23 | *Astroviridae* | + ss RNA | 348 | Yes | Whole genome | 6363 |
| FERRET PESTIVIRUS |  | Ferret | M15 | *Flaviviridae* | + ss RNA | 81 | Yes | Whole genome | 11472 |
| FERRET PESTIVIRUS |  | Ferret | M16 | *Flaviviridae* | + ss RNA | 117 | Yes | Whole genome | 11566 |
| FERRET PESTIVIRUS |  | Ferret | M17 | *Flaviviridae* | + ss RNA | 38 | Yes | Whole genome | 11468 |
| FERRET PESTIVIRUS |  | Ferret | M20 | *Flaviviridae* | + ss RNA | 17 | Yes | Whole genome | 11456 |
| POSSUM HEPACIVIRUS |  | Possum | M23 | *Flaviviridae* | + ss RNA | 3382 | No | Whole genome | 9122 |
| FERRET HEPATITIS E VIRUS |  | Ferret | M14 | *Hepeviridae* | + ss RNA | 391 | No | Whole genome | 8733 |
| FERRET HEPATITIS E VIRUS |  | Ferret | M15 | *Hepeviridae* | + ss RNA | 360 | No | Whole genome | 8737 |
| FERRET HEPATITIS E VIRUS |  | Ferret | M16 | *Hepeviridae* | + ss RNA | 119 | No | Whole genome | 8740 |
| FERRET HEPATITIS E VIRUS |  | Ferret | M17 | *Hepeviridae* | + ss RNA | 1464 | No | Whole genome | 8942 |
| WEASEL JEILONG VIRUS |  | Weasel | M10 | *Paramyxoviridae* | - ssRNA | 1 | Yes | RdRp, partial CDS | 496 |
| HEDGEHOG HEPATOVIRUS |  | Hedgehog | M24 | *Picornaviridae* | + ss RNA | 35817 | No | Whole genome | 7073 |
| FERRET PARECHOVIRUS 2 |  | Ferret | M5 | *Picornaviridae* | + ss RNA | 1.44 | No | Polyprotein, partial CDS | 981 |
| FERRET PARECHOVIRUS 2 |  | Ferret | M16 | *Picornaviridae* | + ss RNA | 0.15 | No | Polyprotein, partial CDS | 326 |
| HUMAN ROTAVIRUS A |  | Hedgehog | M24 | *Sedoreoviridae* | ds DNA | 4.88 | No | RdRp, partial CDS | 379 |

### Double stranded DNA viruses

We identified two non-overlapping adenovirus fragments in ferrets sampled from the North Island, which we assume belong to the same virus (Fig. S1A). This adenovirus belongs to the *Mastadenovirus* genus which we have provisionally named *Ferret mastadenovirus*. This virus was most closely related to *Polar bear adenovirus 1* (25), with 73% amino acid identity. In addition, a short fragment of *Human rotavirus A* from the *Sedoreoviridae* family was found in hedgehogs sampled from the South Island at low abundance (<5 RPM, Fig. S1D, Table 1). The virus could not be genotyped for G or P type due to the short fragment length (379 nucleotides), however the closest known genetic relative was Human rotavirus A strain RVA/Human-tc/JPN/K8/1977/G1P[9] sampled in Japan, with 94% nucleotide sequence similarity. A small number of human transcripts were present in the hedgehog library (accounting for 1.33% of the total read count), thus it is possible that the detected rotavirus fragment originated from trace human contamination rather than active infection of the hedgehog. This level of human contamination was similar to all other libraries (which ranged from 0.75 – 5.33% human reads) where no human-infecting viruses were found.

### Single stranded RNA viruses

We identified an astrovirus in possums across two libraries sampled in the South Island, which we have provisionally named *Possum astrovirus*. A complete viral genome (6,363 nucleotides) as well as a near-complete genome (5,446 nucleotides) were obtained. These sequences were 86% similar at the nucleotide level, and the ORF1b proteins (containing the RNA-dependent RNA polymerase, RdRp) were 94% similar at the amino acid level. We therefore considered these sequences to be from the same virus species. Like other astroviruses, *Possum astrovirus* had an ORF1a (encoding the protease), ORF1b (RdRp) and ORF2 (capsid) (Fig 3B). It also had the ribosomal frame shift (AAAAAAC) near the end of ORF1a, to create ORF1a/b. *Possum astrovirus* was most closely related to *Tasmanian devil associated astrovirus 1* (26), with 56% amino acid sequence similarity. These two viruses formed a clade basal to the classified *Mamastrovirus* genus, but with strong bootstrap support (100%) (Fig. 3A).

**FIG 3.**
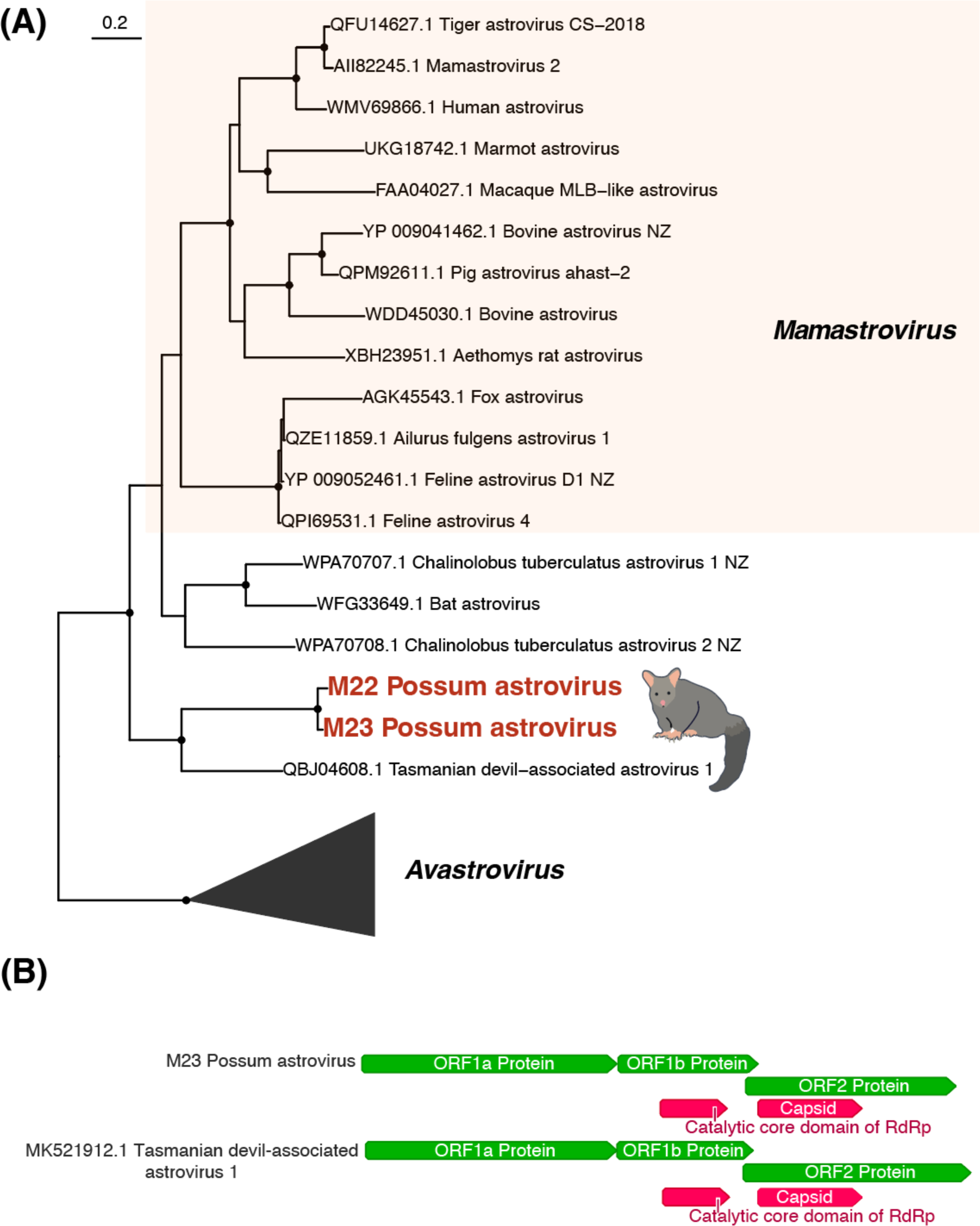
(A) Maximum likelihood phylogenetic tree of the *Astroviridae* (representative viruses only, n = 23 including collapsed clades) based on the ORF1b protein (alignment length of 343 amino acids post-trimming). Viruses identified in this study are shown in red and related viruses are shown in black. Black circles on nodes indicate bootstrap support values of >90%. Branches are scaled according to the number of amino acid substitutions per site, shown in the scale bar. The trees are midpoint rooted for purposes of clarity only. (B) Genome structure of *Possum astrovirus* and the closest known relative (*Tasmanian devil-associated astrovirus 1*).

Five viruses across two viral species were found within the *Flaviviridae* in ferrets and possums. We identified four full genomes of a novel pestivirus in ferrets sampled from across the South Island, now termed *Ferret pestivirus*. These sequences were 93 – 95% similar at the nucleotide level but were highly divergent from other pestiviruses with the closest known virus, *Rhinolophus aFinis pestivirus 1*, sharing only between 42 and 47% amino acid sequence similarity (Fig. 4A). Like other pestiviruses, these genomes consisted of a single polyprotein approximately 11.4kb in length (Fig. 4B). We also identified the whole genome of *Possum hepacivirus* at high abundance (>3000 RPM, Table 1) with 96% amino acid and 85% nucleotide sequence similarity and the same genome structure (Fig. 4C) to the virus first identified in Australian brushtail possums (27).

**FIG 4.**
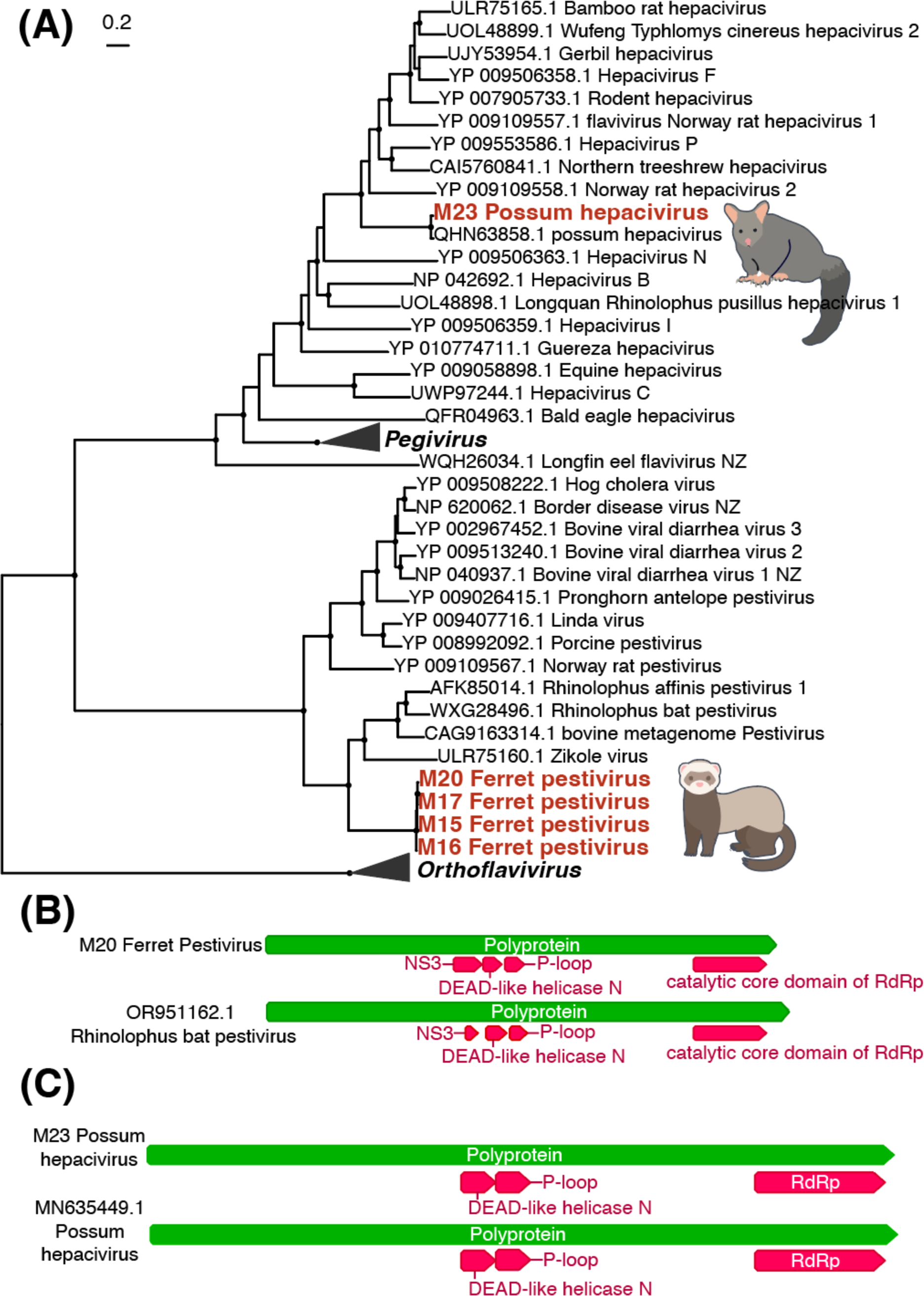
(A) Maximum likelihood phylogenetic tree of the *Flaviviridae* (representative viruses only, n = 47 including collapsed clades) based on the polyprotein (alignment length of 2090 amino acids post-trimming). Viruses identified in this study are shown in red, while related viruses are shown in black. Black circles on nodes show bootstrap support values of >90%. Branches are scaled according to the number of amino acid substitutions per site, shown in the scale bar. The trees are midpoint rooted for clarity only. (B) Genome structure of *Ferret pestivirus* found in this study and a close relative (*Rhinolophus bat pestivirus*). (C) Genome structure of *Possum hepacivirus* found in this study and *Possum hepacivirus* found previously in Australia.

Full genomes of *Ferret hepatitis E virus* were recovered from ferrets sampled from across the South Island, sharing between 92 – 98% amino acid sequence similarity with each other, and up to 92% similar to previously identified strains of this virus (Fig. 5). Phylogenetic analysis of the whole genome showed clear geographic clustering, with the New Zealand strains being most closely related to those sampled in Japan (28) (Fig. 5B). Within New Zealand there appear to be two distinct strains (denoted strain 1 and 2), with 86 – 88% nucleotide sequence similarity between strains, and 95 - 96% nucleotide sequence similarity within the strains. Strain 1 was found in the lower and upper part of the South Island (Southland and Marlborough), while strain 2 was found closer to the centre of the South Island (Canterbury).

**FIG 5.**
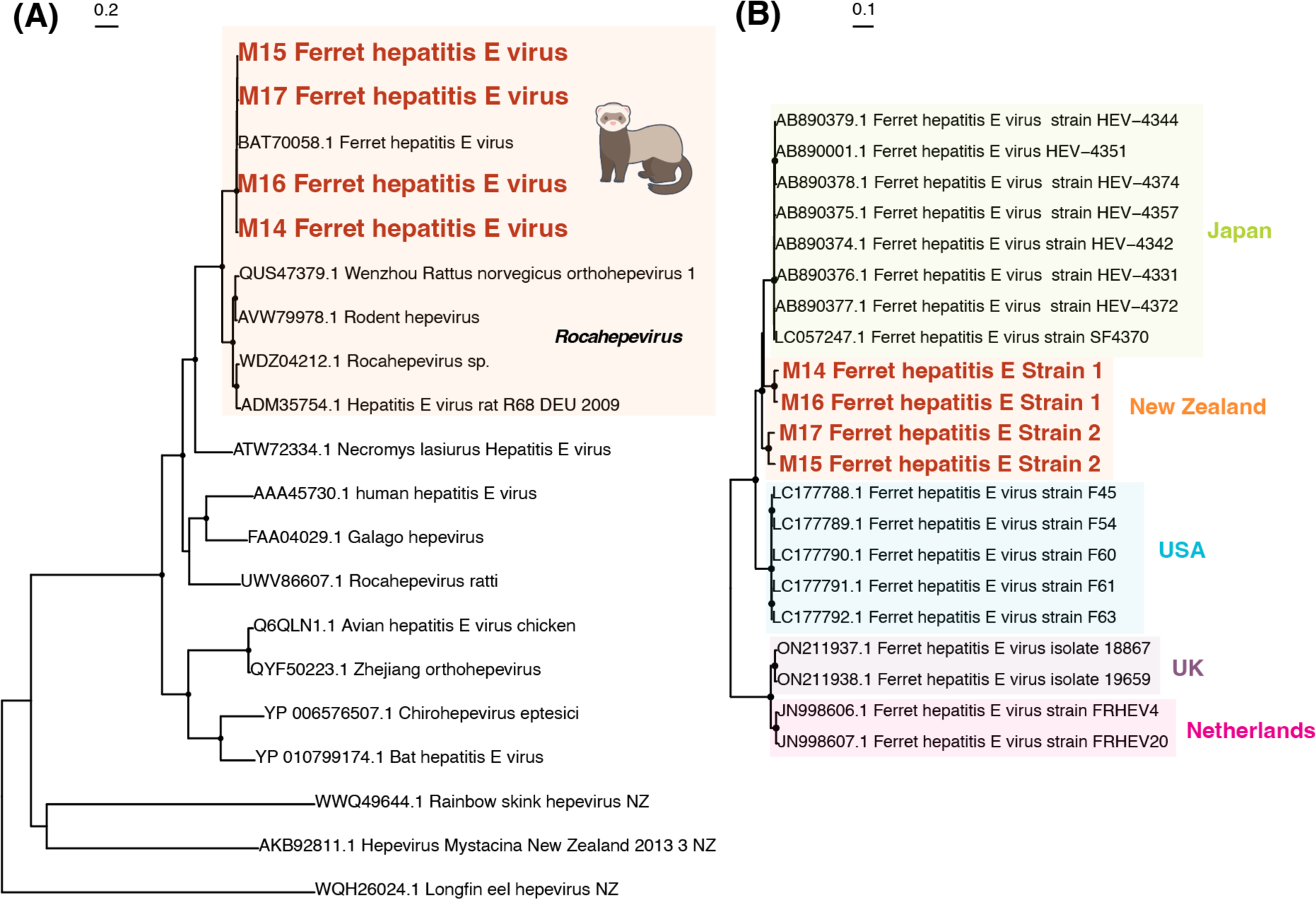
(A) Maximum likelihood phylogenetic tree of the *Hepeviridae* (representative viruses only, n = 20) based on the non-structural polyprotein (alignment length of 1250 amino acids post-trimming). Viruses identified in this study are shown in red, related viruses shown in black. (B) Maximum likelihood phylogenetic tree of *Ferret hepatitis E* strains based on the whole genome at nucleotide level. Shading denotes the country the viruses were sampled. Black circles on nodes show bootstrap support values of >90%. Branches are scaled according to the number of amino acid (A) or nucleotide (B) substitutions per site, shown in the scale bar. The trees are midpoint rooted for purposes of clarity only.

Within the *Arteriviridae*, we identified *Hedgehog arterivirus*, previously found in European hedgehogs in the United Kingdom (29), sharing 92% amino acid sequence similarity (Fig. 6A). We also found two whole genomes of *Wobbly possum disease virus* (WPDV) in possums sampled from the South Island, with 86% nucleotide sequence similarity to each other. These genomes had the same structure as other WPDV sequences, with a large ORF1a/b polyprotein and a frame shift at a ribosomal slippage site (Fig. 6C). Phylogenetic analysis of these viruses along with the seven already identified whole genomes showed that they clustered with WPDV previously identified in New Zealand in 1995, sharing 94 – 96% amino acid with those previously identified (22) (Fig. 6B). The viruses we identified formed a clade with this New Zealand virus, while the WPDV found in Australia formed a sister clade (Fig. 6B).

**FIG 6.**
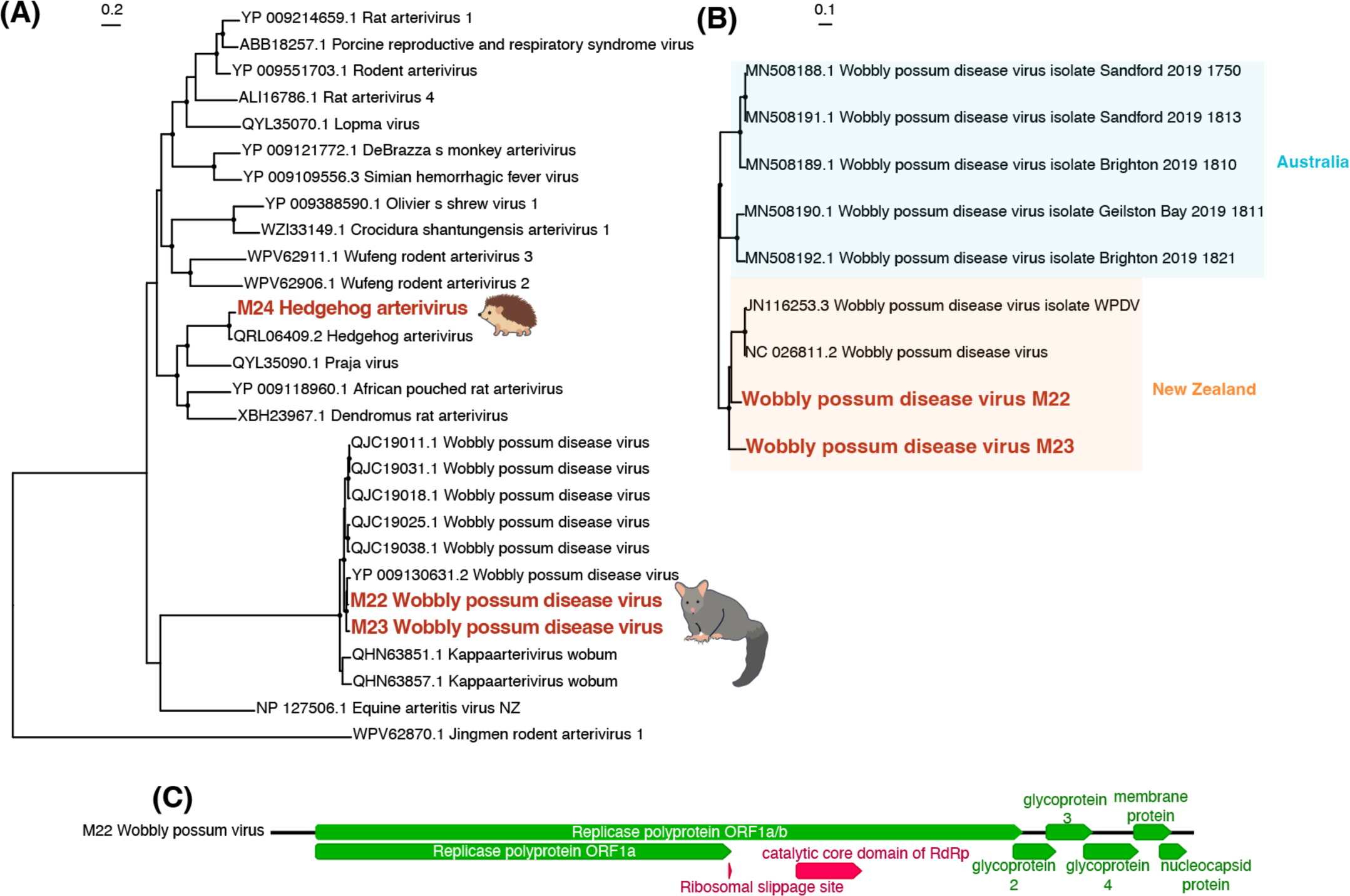
(A) Maximum likelihood phylogenetic tree of the *Arteriviridae* (representative viruses only, n = 28) based on the ORF1a/b polyprotein (alignment length of 1194 amino acids post-trimming). Viruses identified in this study are shown in red, while related viruses shown in black. (B) Maximum likelihood phylogenetic tree of *Wobbly possum disease virus* (WPDV) strains based on the whole genome at nucleotide level. Shading denotes the country in which the viruses were identified. Black circles on nodes show bootstrap support values of >90%. Branches are scaled according to the number of amino acid (A) or nucleotide (B) substitutions per site, shown in the scale bar. The trees are midpoint rooted for purposes of clarity only. (C) Genome structure of the WPDV found in this study.

Two virus species belonging to the *Picornaviridae* were identified in hedgehogs and ferrets. We found *Hedgehog hepatovirus* in very high abundance (>35,000 RPM, Table 1), with 98% amino acid sequence similarity to the virus previously found in European hedgehogs in Germany (30) (Fig. S1C). We also identified a novel parechovirus in two libraries of ferret samples, which we have provisionally named *Ferret Parechovirus 2*. The sequences in the two libraries do not overlap, yet we have conservatively assumed they are the same virus species (Fig. S1C). These viruses are most closely related to *Ferret parechovirus* found in ferrets in the Netherlands (31) with 78 and 88% amino acid sequence similarity, and *Parechovirus sp*. QAPp32 found in the common pipistrelle bat, *Pipistrellus pipistrellus* (80 and 74% amino acid sequence similarity).

Finally, a virus from the genus *Jeilongvirus* in the *Paramyxoviridae* family was found in weasels sampled in the North Island and has been provisionally named *Weasel jeilongvirus*. We recovered three non-overlapping fragments of this virus, which we have conservatively assumed to be the same virus (Fig. S1B). These fragments are most closely related to *Feline paramyxovirus 163* found in Japan (84 – 89%).

## DISCUSSION

This study presents the first metatranscriptomic survey of viruses in five introduced mammalian species in New Zealand: ferrets, stoats, weasels, brushtail possums and hedgehogs. We identified 11 mammalian-infecting viral species spanning eight viral families, including four novel virus species within the *Adenoviridae, Astroviridae, Flaviviridae* and *Paramyxoviridae*. Viral abundance varied considerably across species, with the highest viral load observed in hedgehogs (driven by *Hedgehog hepatovirus*), followed by possums and select samples of ferrets. Notably, complete genomes were recovered for six viruses, underscoring the utility of metatranscriptomics for in-depth viral discovery. These findings expand our understanding of the mammalian virome in New Zealand and highlight previously unrecognized viral diversity in introduced hosts.

Four novel viruses were identified in this study: *Ferret mastadenovirus*, *Possum astrovirus*, *Ferret pestivirus* and *Weasel jeilongvirus*. These viruses were phylogenetically distinct from known genetic relatives and, in some cases, highly divergent showing less than 60% amino acid sequence similarity, uncovering previously uncharacterized viral lineages. Genetic relatives of these viruses include known animal and zoonotic pathogens, such as Human astrovirus VA1/HMO-C associated with gastroenteritis, and closely related to bovine and mink astroviruses (32). While no clear zoonotic threats were identified in our data, these findings provide useful surveillance for public and animal health, particularly given the ecological overlap of these mammalian hosts and their proximity to humans, native wildlife and domestic animals.

Several viruses identified in this study have been previously described, including WPDV, *Possum hepacivirus*, *Ferret hepatitis E virus* and *Hedgehog arterivirus*, all sampled in animals from around the world (27, 33, 34). The detection of these viruses in New Zealand suggests long-term persistence following a single introduction along with their hosts in the 19^th^ and 20^th^ centuries. Population bottlenecks caused by small founding population sizes (35–37) could have eliminated many viruses from these mammal populations at the time of their introduction to New Zealand, but our results indicate that at least some viruses were introduced to New Zealand along with their hosts. Notably, we also identified *Human rotavirus A* in a hedgehog sample, which raises the possibility of reverse zoonotic transmission, likely via environmental contamination. To our knowledge, *Human rotavirus A* has not been found in hedgehogs previously, however hedgehogs are increasingly being recognized as potential reservoirs for zoonotic viruses (38), and their presence in urban or peri-urban environments could facilitate such spillovers. Human rotaviruses have occasionally been detected in non-human animals including domestic animals and non-human primates (39, 40), lending weight to the possibility that reverse zoonotic transmission has also occurred in this case. However, it is also possible that the *Human rotavirus A* detected in the hedgehog library reflects contamination during sample collection or processing, rather than true infection. The nature of our sampling method (sourcing animals from established trapping programmes) meant that human contamination was unavoidable. Further sampling of hedgehogs coupled with targeted PCR, in situ hybridization, or serology would be necessary to determine whether active infection occurred.

The phylogenetic patterns observed in both WPDV and *Ferret hepatitis E virus* revealed clear geographic clustering of New Zealand viruses. The WPDV genomes identified in this study clustered with the previously described New Zealand strain (22), forming a distinct clade sister to Australian WPDV lineages (27). This pattern is consistent with historical records of possum introductions, which occurred only in the early 1900s (11), and indicates that WPDV has likely been circulating and evolving within New Zealand possum populations since. Similarly, *Ferret hepatitis E virus* from New Zealand formed a well-supported clade most closely related to Japanese strains (28), suggesting a shared origin or common ancestral introduction. The distinct separation of these viruses by geographic region reflects host movement restrictions and ecological isolation, as well as evolutionary divergence following limited introduction events. Such patterns underscore the importance of localized viral surveillance and suggest that introduced mammalian species in New Zealand may harbour uniquely evolved viral lineages with potential implications for both animal health and biosecurity. These patterns also suggest that the strong biosecurity measures in New Zealand have successfully prevented further introductions of these mammals from overseas.

Despite the close evolutionary relationship and geographic overlap among mustelid species (24), we found no evidence of viral sharing between hosts. This result is surprising given the potential for cross-species viral transmission, particularly among mustelids (ferrets, stoats and weasels). Evidence for spillover events of *Aleutian Mink Disease Virus* from mink (*Neogale vision*) was found in six other mustelid species in Poland (41) and eight in Finland (42), demonstrating the ability of some viruses to seemingly jump between mustelid species. The lack of evidence for cross-species viral transmission in our study may be an artifact of our opportunistic sampling, which included relatively few individuals from stoats and weasels compared to ferrets, reducing the probability of detecting low-frequency transmission events or shared virus species. Additionally, differences in habitat use, (e.g. ecological niche separation (43)), behaviour (solitary lifestyle (44)) or immune responses could contribute to barriers to viral exchange (45, 46). More extensive and systematic sampling across all species and regions is required to fully assess the potential for viral spillover between these hosts.

New Zealand has the ambitious goal of becoming free of mammalian predators by 2050 (47). Biological control of pest animals, such as those sampled in this study, could be a key tool for reaching this goal (48, 49). Indeed, the discovery of host-specific viruses highlights intriguing possibilities for biological control. Nevertheless, for effective biocontrol, viruses need to be highly pathogenic, species-specific, and safe for non-target species - including humans, native wildlife and domestic animals. All of the viruses we detected, including the novel pestivirus in ferrets and astrovirus in possums, were identified in apparently healthy individuals where no overt signs of disease were noted. The exception may be WPDV, which has been associated with neurological disease in brushtail possums (27), as well as *Hedgehog arterivirus*, associated with neurological disease and fatal encephalitis (33, 50), although as above no animals sampled had obvious signs of disease. Rigorous assessment of pathogenicity and species specificity is required prior to any future application. More broadly, the use of viruses as biocontrol agents remains contentious (51) and would require robust risk assessment and regulatory oversight, especially in a biodiverse and conservation-sensitive context like New Zealand.

Our findings highlight several promising avenues for future research. First, increasing the sample size and geographic coverage for underrepresented species such as stoats, weasels and hedgehogs would allow for a more robust assessment of viral diversity and potential cross-species viral transmission. Second, functional studies of the novel viruses identified here could help elucidate their host range, transmission potential and pathogenicity. Third, integrating ecological data such as species movement, diet and habitat overlap could shed light on the mechanisms shaping viral communities. Finally, monitoring for known and novel viruses in native New Zealand wildlife is essential to assess the potential for spillover from introduced species, particularly as efforts to manage or eradicate invasive mammals continue. Together, these findings underscore the value of viral surveillance in introduced wildlife and provide a crucial foundation for understanding and mitigating emerging infectious disease risks in New Zealand’s unique ecosystem.

## MATERIALS AND METHODS

### Sampling and RNA extraction

Kidney and liver samples were collected from 290 individuals from five introduced mammalian species: ferrets (*Mustela furo*), stoats (*Mustela erminea*), weasels (*Mustela nivalis*), brushtail possums (*Trichosurus vulpecula*) and European hedgehogs (*Erinaceus europaeus*), from the upper North Island and South Island of New Zealand in 2021 (Fig. 1). Animals were either live trapped and culled, kill trapped or opportunistically collected as fresh roadkill. All animals that were trapped and killed were done so as part of already established pest control efforts. Upon collection, animals were frozen at −20°C and transferred to the University of Auckland (for North Island samples) or the University of Otago (for South Island samples). Before dissection, carcasses were thawed and checked for signs of decomposition to determine suitability for RNA extraction. Liver and kidney tissue samples were harvested and stored in RNALater (Thermo Fisher Scientific) at −80°C until RNA extraction using the Qiagen RNeasy Plus Mini kit (Qiagen) following the manufacturer’s instructions. RNA from up to 30 individuals from each species was pooled per location and ribosomal RNA was removed using the Ribo-Zero-Gold Kit from Illumina, then sequenced using Illumina NovaSeq 6000 at the Australian Genome Research Facility (AGRF), Melbourne, Australia.

### Viral discovery

Sequencing reads were quality-trimmed using Trimmomatic (v0.38), with removal of adapter sequences and bases with a quality score below 5 using a sliding window of four bases (52). Additionally, low-quality bases (quality score <3) were trimmed from the ends of reads, and sequences shorter than 25 nucleotides were discarded. Following quality control, reads were *de novo* assembled into contigs using MEGAHIT (v1.2.9) (53). These contigs were then compared to the NCBI nucleotide (nt) and non-redundant protein (nr) databases using BLASTn (BLAST+ v2.13.0, (54)) and DIAMOND (DIAMOND v2.1.6, (55)) to identify viral sequences. To reduce false positives, similarity thresholds of 1 × 10⁻⁵ for the nt database and 1 × 10⁻¹⁰ for the nr database were applied. Virus abundance was estimated by mapping reads back to the assembled contigs using Bowtie2 (v2.4.5, (56)) and SAMtools (v1.9, (57)). Viral sequences present at read counts <0.1% of those in another library and >99% identical at the nucleotide level were considered likely cross-contamination due to index hopping and excluded from further analysis. The probable host origin of each virus was inferred based on phylogenetic relatedness to known viruses. Viruses that clustered with known mammalian viruses were subject to further evolutionary analysis. Viruses that were phylogenetically distinct from vertebrate host viruses were assumed to be more likely associated with diet, microbiome or environmental sources.

### Phylogenetic analysis

Putative viral transcripts were first translated and then aligned with representative protein viral sequences from the same viral genus or family retrieved from GenBank. These representative sequences were chosen to capture diversity within each viral genus or family by selecting a small number of sequences from major clades and including all sequences closely related to the viruses identified in this study. Alignments were performed using MAFFT (v7.402) (58) with the E-INS-i or L-INS-i algorithms (Table S1) and subsequently trimmed using TrimAl (v1.4.1) (59). Phylogenetic trees for each viral family or genus were estimated using the maximum likelihood method in IQ-TREE (v1.6.12), with the program determining the best-fit substitution model using ModelFinder (60) and node robustness evaluated through the approximate likelihood ratio test with 1,000 replicates (61). Sequences found within the same library were considered to represent different viral species (rather than intra-species viral diversity) if overlapping aligned sequences had <90% amino acid identity. Phylogenetic trees were visualized using APE (v5.4) (62) and ggtree (v2.4.1) (63) in R (v4.0.5) (64).

Select whole viral genomes were also further analysed at the nucleotide level. Annotations were conducted using BLAST and the conserved domains database (65). Phylogenetic analysis was conducted at the nucleotide level using whole genome sequences, aligning the sequences using MAFFT. Phylogenetic trees were estimated using the same methods as described above.

## Data availability

The sequence data generated in this study has been deposited in the Sequence Read Archive (SRA) under the project accession number PRJNA1304147. Virus consensus sequences have been deposited NCBI/GenBank with assigned accession numbers XXX.

## ACKNOWLEDGMENTS

JLG is funded by a New Zealand Royal Society Rutherford Discovery Fellowship (RDF-20-UOO-007), the Webster Family Chair in Viral Pathogenesis and a University of Otago Research Grant. RKF was funded by The Australia & Pacific Science Foundation (APSF24008) and a University of Otago Research Grant. FP was funded by a fellowship from New Zealand Predator-Free 2025 (Genomic applications for invasive species control focussing on mustelids-3723891).

Thanks to Vibha Thakur for lab assistance and Leigh Ellmers (Landcare Research Lincoln) for sample contribution. Thanks especially to the many volunteers providing samples and/or giving FP access to their land. To name a few: Predator Free Franklin, Andy Saunders, Willow van Heugten, Karen Tricklebank and Emily Lane.

